# Functional dissection of the *ARGONAUTE7* promoter

**DOI:** 10.1101/392910

**Authors:** J. Steen Hoyer, Jose L. Pruneda-Paz, Ghislain Breton, Mariah A. Hassert, Emily E. Holcomb, Halley Fowler, Kaylyn M. Bauer, Jacob Mreen, Steve A. Kay, James C. Carrington

**Affiliations:** Donald Danforth Plant Science Center, St. Louis, MO, USA; Computational and systems biology program, Washington University in St. Louis, MO, USA; Division of Biological Sciences and Center for Chronobiology, University of California San Diego, La Jolla, CA, USA; Department of Integrative Biology and Pharmacology, McGovern Medical School, Houston, TX, USA; Department of Neurology, University of Southern California, Los Angeles, CA, USA

**Author notes:** Corresponding author: James C. Carrington.

## Abstract

ARGONAUTES are the central effector proteins of RNA silencing which bind target transcripts in a small RNA-guided manner. *Arabidopsis thaliana* has ten *ARGONAUTE* (AGO) genes, with specialized roles in RNA-directed DNA methylation, post-transcriptional gene silencing, and antiviral defense. To better understand specialization among *AGO* genes at the level of transcriptional regulation we tested a library of 1497 transcription factors for binding to the promoters of *AGO1, AGO10*, and *AGO7* using yeast 1-hybrid assays. A ranked list of candidate DNA-binding TFs revealed binding of the *AGO7* promoter by a number of proteins in two families: the miR156-regulated SPL family and the miR319-regulated TCP family, both of which have roles in developmental timing and leaf morphology. Possible functions for SPL and TCP binding are unclear: we showed that these binding sites are not required for the polar expression pattern of *AGO7*, nor for the function of *AGO7* in leaf shape. Normal *AGO7* transcription levels and function appear to depend instead on an adjacent 124-bp region. Progress in understanding the structure of this promoter may aid efforts to understand how the conserved AGO7-triggered *TAS3* pathway functions in timing and polarity.

## 1 Introduction

Small RNAs regulate developmental timing and morphogenesis in a wide range of eukaryotes. Heterochronic (abnormal timing) mutants of the model nematode *Caenorhabditis elegans* led to the discovery of the first microRNA (miRNA)-target pair [1, 2]. Similar screens for *A. thaliana* heterochronic mutants led to elucidation of a specialized pathway in which *trans*-acting small interfering (tasi)RNA are produced from noncoding *TAS3* transcripts [3–6]. Genetic analysis of leaf morphology has also led to the discovery of several other aspects of RNA silencing, including the cloning of the first *ARGONAUTE (AGO)* gene [7]. AGO proteins bind small RNAs and effect small-RNA-guided regulatory changes. Several families of *MIRNA* genes are conserved in all land plants [8], and miRNA from the majority of these families repress TFs controlling developmental programs, suggesting that AGO-miRNA-TF circuits became embedded in the core regulatory networks for the plant body plant early in land plant evolution [9].

The *A. thaliana* genome contains ten *AGO* genes, which function in development, stress resistance, and defense against viruses and transposons [10]. AGO7 and AGO10 are highly specialized: each has limited adaxial and vascular expression [11, 12] and a single main binding partner: miR390 and miR166, respectively [13, 14]. AGO7 triggers production of phased siRNAs from *TAS3* noncoding transcripts [13, 15–17]. Effects on ARF3, ARF4, and possibly ARF2 are the main downstream output of the AGO7/TAS3/SGS3/RDR6/DCL4 pathway [18–21]. AGO7 action is thought to limit production of *TAS3* tasiRNAs such that tasiRNA movement creates a graded accumulation pattern in developing leaf primordia [12, 22]. This gradient contributes to patterning of *ARF* target mRNA, establishing either an opposing gradient or a sharp boundary, which may contribute to robust maintenance of polarity [23]. The *TAS3* pathway has important roles in leaf development in all plants examined thus far, including moss [24], maize [25–27], tomato [28], lotus [29] and alfalfa [30].

Understanding the functions of miRNA such as miR390 and miR166 will require information on the signals controlling tissue-specificity of their AGO partners. Our objective in this work was to identify upstream regulators of *AGO* genes and link them to existing genetic knowledge. We capitalized on new yeast-based tools that provide a fast way to identify upstream regulators. We identified unexpected connections to two other conserved miRNA-TF circuits that control leaf morphogenesis and defined two other functional regions of the *AGO7* promoter.

## 2 Results

### 2.1 Multiple SPLs and TCPs bind the *AGO7* promoter

We sought to identify TFs controlling the expression of the three main *AGO* genes involved in post-transcriptional control of development *(AGO1, AGO10*, and *AGO7)* using high-throughput yeast 1-hybrid assays. Our automated strategy, described previously [31, 32], uses a large collection of arrayed A. *thaliana* TFs (details below) and also short promoter bait sequences, for high resolution and sensitivity. We considered four fragments for each promoter, with ∼50 bp of overlap between fragments, to ensure that fragment-edge binding sites were assayed. For *AGO7* these fragments spanned a 1934 bp region (Figure 1A). Transgenes driven by the collective sequences represented by these fragments are sufficient to complement corresponding *ago* mutants [13, 33, 34], suggesting that they contain the most important upstream regulatory elements. Promoter fragments were screened against a TF-activation domain fusion library in 384-well format with one prey TF per well [31], using *β*-galactosidase reporter activity from fusion to promoterless *uidA* coding sequence as a quantitative readout (Figure 1).

**Figure 1.**
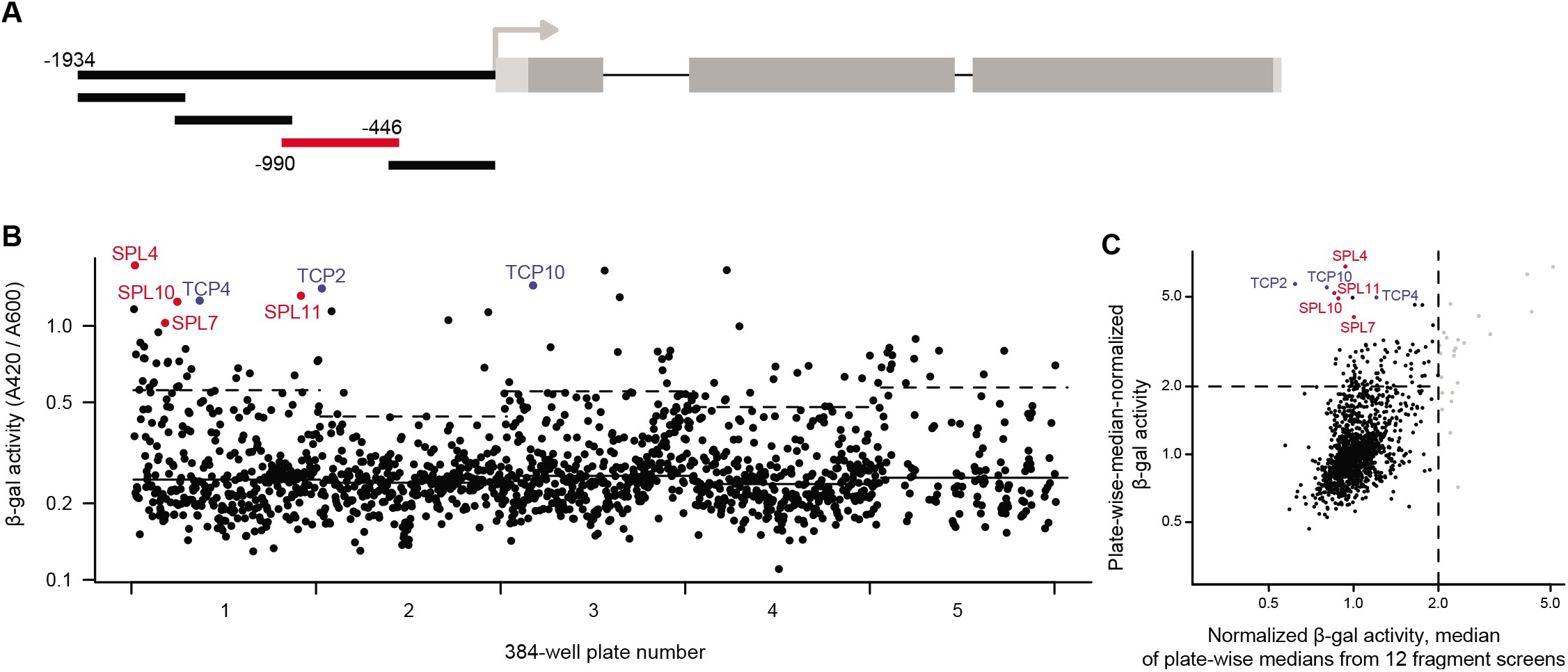
SPL and TCP TFs bind the *AGO7* promoter in yeast. A. Schematic of *AGO7* promoter illustrating four fragments screened with Y1H assays. Subsequent panels show results for the fragment indicated in red, which spans the region from 990 bp to 446 bp upstream of the transcription start site. B. Scatterplot of *β*-gal activities for each prey TF constructs screened. Wells are shown in row-first order for each of the five plates. Median activity for each plate is indicated with solid lines. Dashed lines indicate a cutoff of 6 median absolute deviations above the median for each plate. Hits from SPL and TCP families are highlighted. C. Diagnostic plot incorporating data from 12 screens. Y-dimension reflects the same values as panel B, normalized by plate median. X-dimension results from taking the median of plate-wise-median activities from all twelve *AGO* promoter fragment screens. Vertical dashed line demarcates TFs for which median reporter activity is two-fold higher than the median for their plate (nonspecific activators, light gray).

A total of 1497 TFs were tested for *AGO* promoter binding. This collection consisted mainly of sequence-specific TFs, but also includes transcriptional co-factors and empty vector control wells [31]. Each TF was tested against each promoter fragment a single time. We ranked TF candidates based on normalizing promoter-fragment-driven *β*-gal activity by the median value for each plate (as illustrated in Figure 1B), to account for systematic differences between plates. We separately plotted signal distributions across all twelve screens (Figures S1, S2, and S3) to assess which TFs “hits” act as nonspecific activators in this system, as described below. (See supplemental dataset [35] and summary tables [36] for additional details.)

Of the TFs families assayed, only two were represented by multiple hits 6 median absolute deviations or more above the median for their plate (Figure 1B). The first group, Teosinte Branched/Cycloidea/PCF family factors (TCPs), had previously been suggested to directly regulate *AGO7* [37]. The three TCP hits identified are miR319 targets [38] and redundantly control leaf margin development and senescence [39]. The second group, SQUAMOSA-PROMOTER-BINDING PROTEIN-LIKE (SPL) factors, are master regulators of heteroblasty in A. *thaliana* and other plants [40], the same context in which *AGO7* was discovered [3]. *AGO7* was prioritized over *AGO10* and *AGO10* for mutagenesis and functional analysis based on interest in these TFs, for which roles in timing but not polarity are well-established.

We examined the distribution of reporter activity for other promoter fragments screened, confirming that these SPL and TCPs specifically hit the second proximal region of the *AGO7* promoter. Plate-wise median *β*-gal activities for the SPL and TCP hits were close to the median (across all twelve screens) for their plate (Figure 1C), indicating that they do not fall in the group of TFs that are nonspecific reporter gene activators.

We further tested a group of SPL and TCP factors with a second Y1H system, based on a secreted luciferase reporter with an improved dynamic range [41]; repeated testing reduces statistical false positives and use of alternative reporters can reveal reporter-gene-specific technical false positives [42]. This secondary screening confirmed that multiple SPL and TCP TFs bind the second proximal *AGO7* promoter fragment tested, despite considerable experimental variability (Figure S4). Some TFs yielded a small degree of activation relative to two different empty vector controls; it is not clear whether these small differences reflect lack of binding (i.e. nonspecific binding only) or indicate binding that is weak but specific.

We assessed possible SPLs and TCPs binding sites using DNA-binding specificity models determined based on *in vitro* sequence affinity with protein-binding microarrays [43]. These position-weight matrices (PWM, downloaded from the CisBP database) match consensus binding sequences previously determined with *in vitro* selection for SPLs [44] and TCP4 [39]. These targeted scans complement a wider analysis done with the ‘Find Individual Occurences of Motifs’ tool (FIMO) tool [45] and three collections of *A. thaliana* TF PWMs [43, 46, 47] documented in the supplemental materials [35]. An example sequence logo for one model, for SPL11, is shown in Figure 2A. Because the Y1H bait of interest extends to position −990 (Figure 1A), we considered the 1 kb region adjacent to the annotated *AGO7* transcription start site. For SPL11, the highest-scoring positions (on both strands) were centered on the only two ‘GTAC’ motifs (SPL core binding sites) in that region, at −500/−497 and −486/−483 (Figure 2, panels B and C).

**Figure 2.**
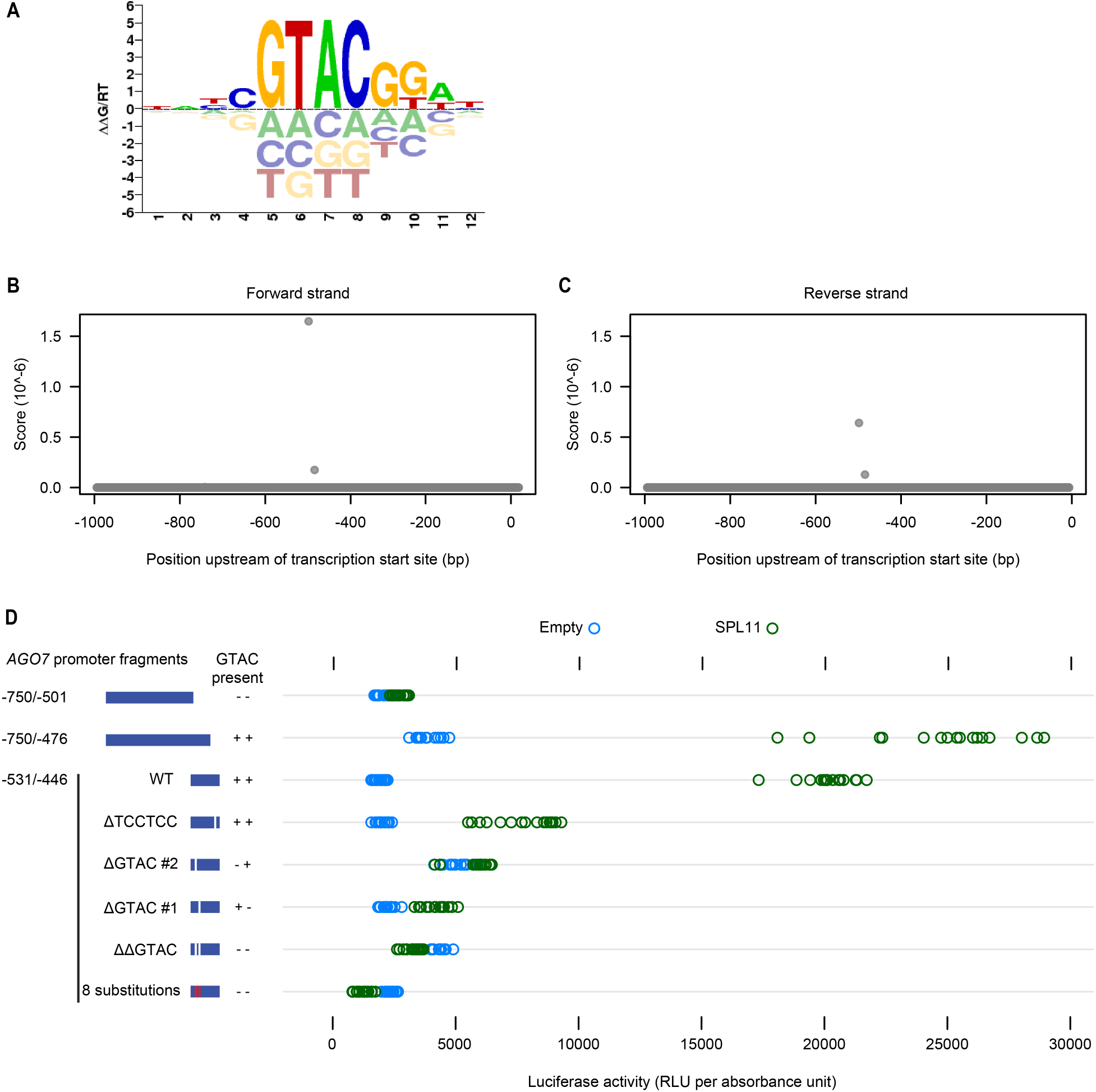
Identification of SPL11 binding sites. A. Sequence logo for SPL11 PWM, as downloaded from CisBP. Individual position weights can be interpreted as binding specificity contributions (changes in free energy, arbitrary units). B and C. Scores for SPL11 PWM at each position of the 1 kb region upstream of the annotated *AGO7* transcription start site. D. Reporter activity (relative luminescence units normalized by A600) for SPL11 and pDEST22 (empty vector) tested in yeast against *AGO7* promoter baits including several derivatives of the −531/−446 region. Modifications included one or two 4-bp deletions, 8 substitutions (TCCG/AAGG), and an unrelated 6-bp deletion; see Table 2, below.

We tested the significance of these core ‘GTAC’ sequences using the luciferase reporter gene in yeast. Truncated Y1H bait sequences (−531/−446 and −750/−476) containing core binding sequences yielded activation of the reporter when tested against SPL11, but not with the corresponding empty prey vector (Figure 2D). By contrast, activation was not observed for a 3’-truncated bait lacking ‘GTAC’ sites (−750/−501), nor for modified −531/−446 bait sequences with one or both 4-mers deleted or scrambled (Figure 2D). Deletion of an unrelated 6-bp region reduced reporter activation (compared to empty vector) but not to the same extent. These results are consistent with direct SPL binding, possibly with some degree of cooperativity, at one or both ‘GTAC’ sites in the yeast system.

We similarly scanned the promoter sequence with empirically determined PWM for five of eight *CINCINNATA-like* TCPs, a set that includes four of the five miR319 targets in *A. thaliana* [38, 48] but does not include TCP10. The highest scoring positions for four TCPs were centered on a ‘TGGTCC’ motif at −459/−454 (Figure 3, panels E to I). This 6-mer was the most highly enriched sequence in the promoters of a set of experimentally defined TCP targets [39], and is present in the “most preferred” sequences for TCP3, TCP4, and TCP5 PWMs. A second ‘TGGTCC’ site at −428/−423 was among the four highest-scoring sequences for all five TCPs considered (Figure 3), but was absent from the −990/−446 region that yielded TCP hits in the initial Y1H screen. High-scoring positions for the TCP2 PWM included a related ‘GGGACC’ sequence at −764/−770 followed by the −459/−454 ‘TGGTCC’ motif (Figure 3, panels A and F). The second highest scoring position for TCP24 was centered on a nearby ‘GTTCCC’ sequence (Figure 3J).

**Figure 3.**
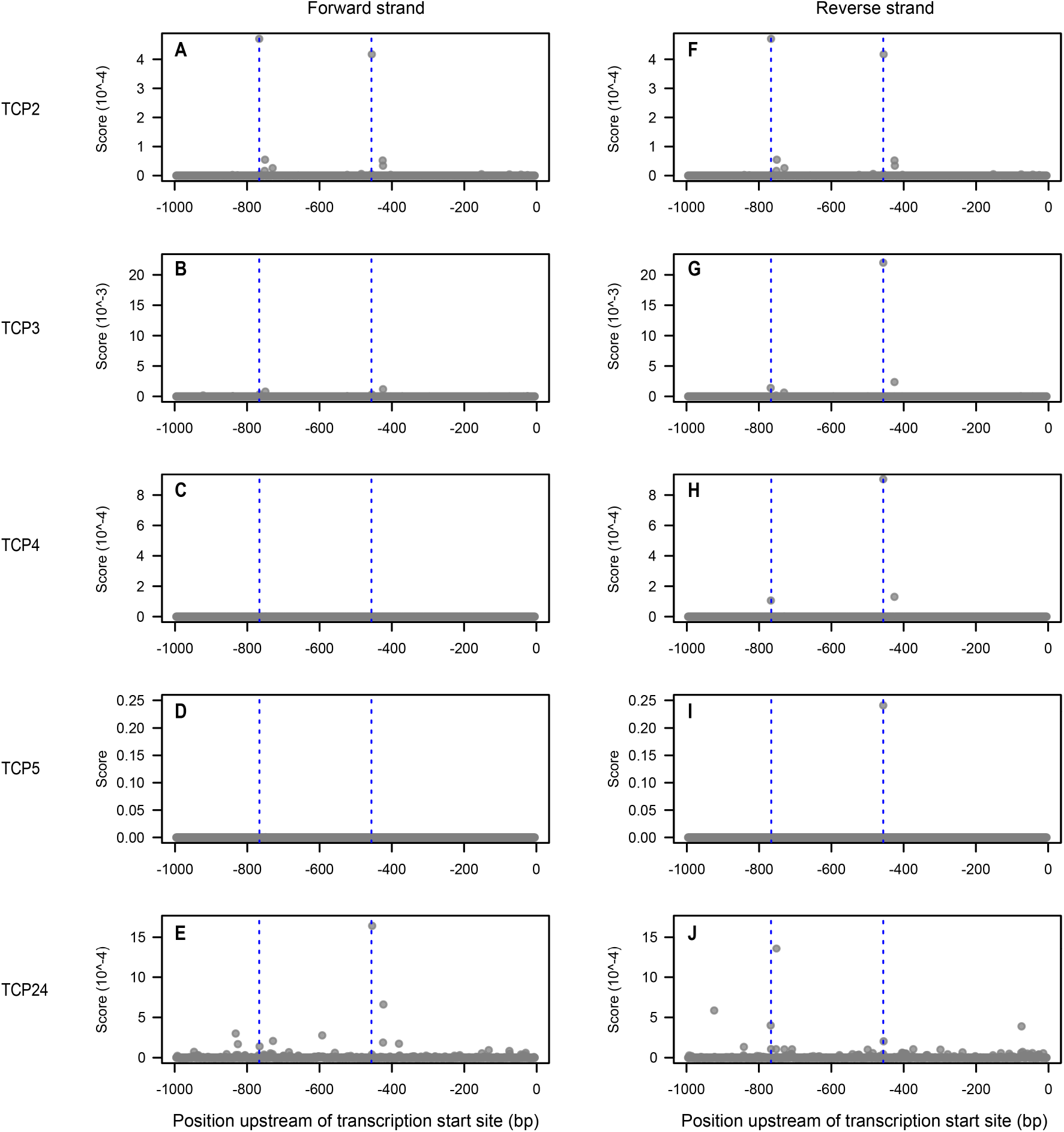
Identification of TCP binding sites. A to J. PWM scores at each position of the 1 kb region upstream of the annotated *AGO7* transcription start site for the TCPs indicated. Dashed blue lines indicate the two highest scoring positions for TCP2: a ‘GGGACC’ sequence at −764/−770 and a ‘TGGTCC’ sequence at −459/−454. Panels A and F are identical, because the CisBP model for TCP2 is perfectly symmetrical.

We tested requirements for candidate TCP binding sites with the luciferase Y1H system. Truncated bait sequences (−750/−501 and −750/−476) lacking all four sites described above did not drive reporter activation (relative to the empty prey vector control) when tested with TCP2 (Figure 4). The −990/−446 region used in the initial screen yielded reporter induction, as did a 5’-truncated 86 bp bait region (−531/−446) containing the higher-scoring ‘TGGTCC’ motif (Figure 4). The same truncated bait sequence with the ‘TGGTCC’ 6-mer deleted did not yield reporter activation (Figure 4). We conclude that the −459/−454 ‘TGGTCC’ is a high-affinity TCP binding site that functions in the yeast system and possibly *in planta*.

**Figure 4.**
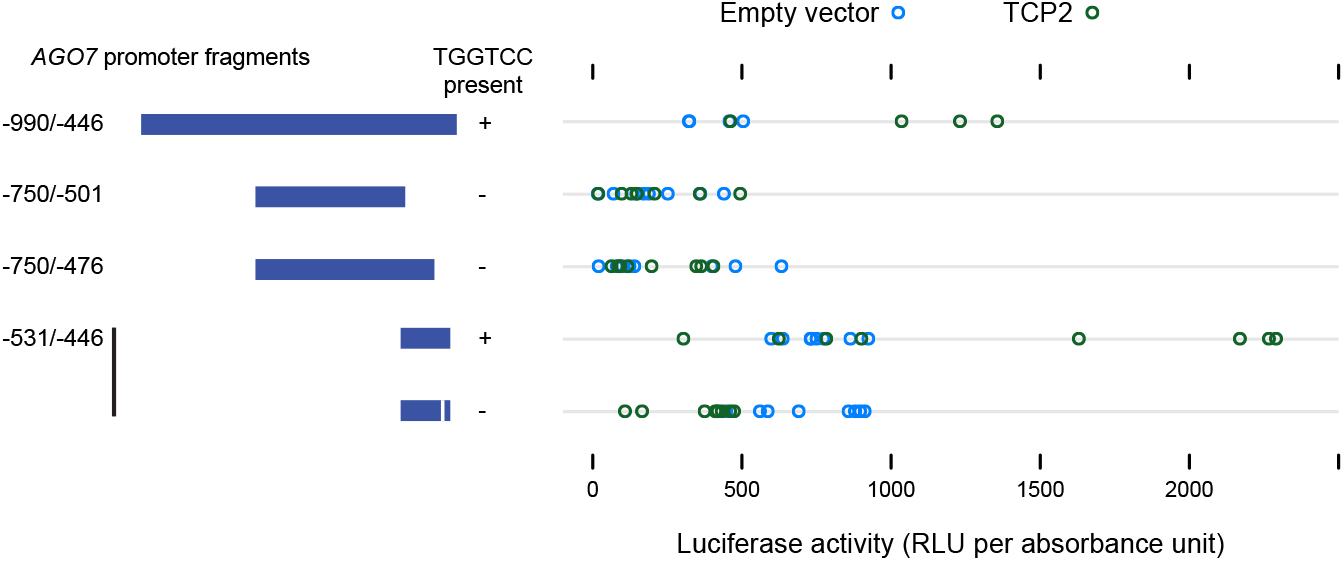
Testing of TCP binding sites in yeast. Reporter activity for TCP2 and pDEST22 (empty vector) tested against Y1H bait from initial screens (−990/−446), two truncated versions lacking candidate TCP binding sites, and the −531/−446 region, with candidate TCP binding site (‘TGGTCC’) deleted or intact.

### 2.2 SPL binding sites are not required for polar *AGO7* transcription

To test the possibility that SPL and/or TCP binding sites contribute to polar *AGO7* transcription, we fused a series of truncated versions of the *AGO7* promoter to *GUS* for comparison to previously described transcriptional reporter lines [12, 13]. Consistent with previous results [12], the 1934 bp region upstream of the annotated *AGO7* transcription start site yielded clear adaxial signal in transverse sections of leaf primordia (Figure 5A). A 482 bp version of the promoter yielded the same pattern in almost all plants tested (Figure 5B), indicating that SPL core binding sites (−500/−496 and −486/−483) are not required for this pattern. By contrast, the TSS-proximal 298 bp region rarely yielded visible blue reporter signal (Figure 5C). Weak adaxial signal was visible for a small proportion of plants, including 2 of 7 plants for one of two transgenic families for the experiment illustrated (Figure 5C; see also the supplementary photomicrograph dataset [49]). Given that promoterless-GUS transformants did not yield visible blue signal (Figure 5D) in any of our experiments, this raises the possibility that *cis* elements in the proximal 298 bp region can confer adaxial polarity to *AGO7* transcription. The −482/−299 region, however, is a larger determinant of *AGO7* transcription level, as discussed further below.

**Figure 5.**
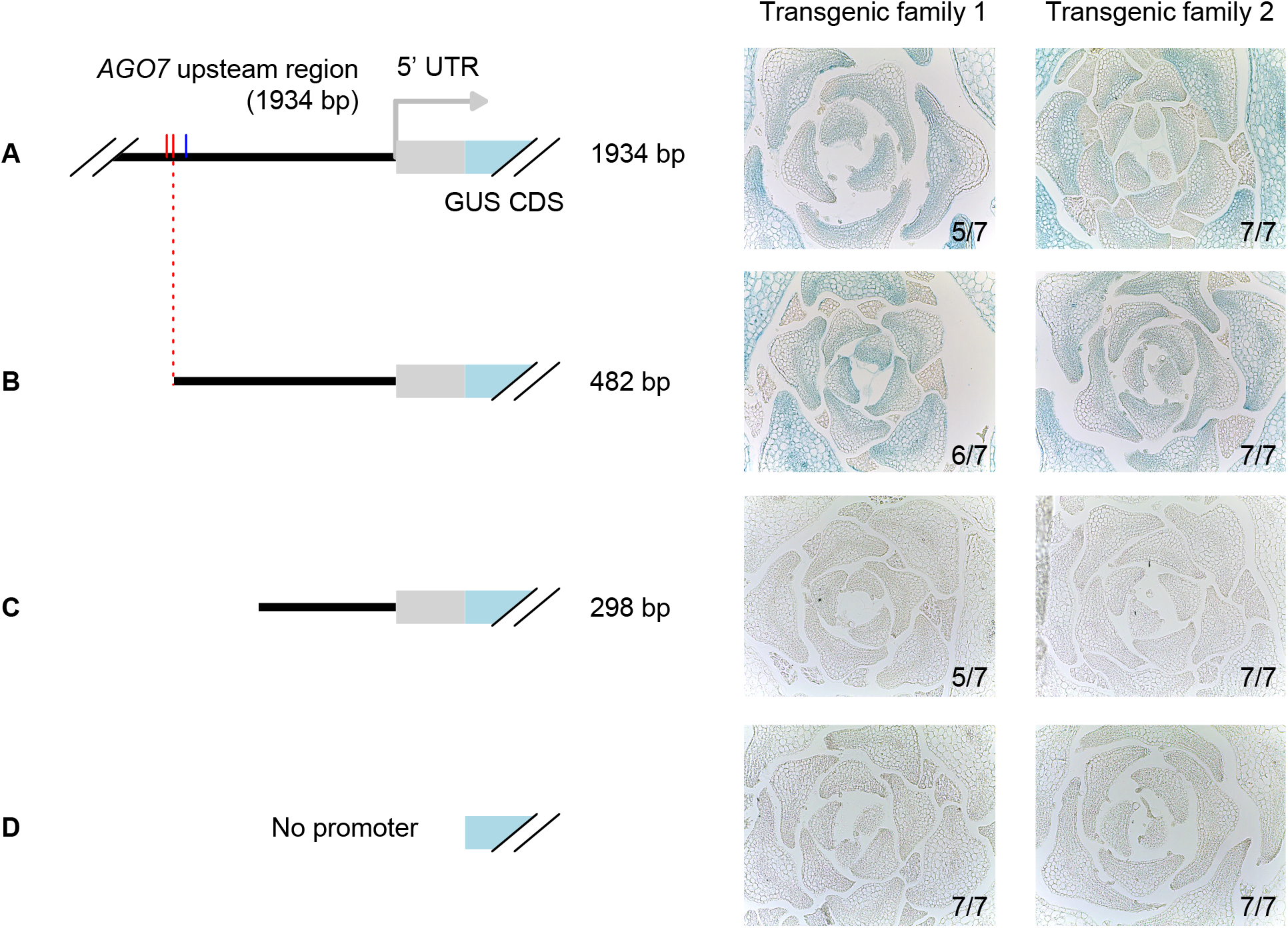
Histological analysis of GUS reporter gene activity driven by truncated *AGO7* promoter constructs. Core SPL binding sites are indicated in red; the 482 bp promoter construct illustrated in panel B ends immediately adjacent to the second site. Blue tick mark indicates TCP binding motif at −459/−454. For each construct, results are shown for two independent transgenic families (groups of T_3_ siblings, each descended from a different transformant; each group was stained in a separate scintillation vial). The predominant class for each family (categorized as having distinct abaxial signal or not) is illustrated with a representative transverse section through young leaf primordia. The number of plants in the majority class is indicated as a fraction. Two and one plants yielded a weaker and/or less strongly adaxial pattern than shown here for 1934 bp and 482 bp promoter constructs, respectively. Between three and nineteen independent lines were tested for all of the constructs shown here, with broadly similar results across multiple experiments.

### 2.3 SPL and TCP binding sites are not strictly required for *AGO7* function

We similarly tested *cis* requirements for transgenic complementation of *ago7* mutants. We inserted a series of truncated versions of the *AGO7* promoter upstream of the *AGO7* coding sequence (including an N-terminal 3x-hemagglutinin (HA) tag). Previous results [13] indicated that the 1934 bp promoter version of this transgene is functional for complementation of transformed *ago7* mutants. For the experiment illustrated in Figure 7, blinded classification of downward leaf curling assigned 100% of empty-vector-transformed reference genotype plants *(ago7* mutant and wild-type Col-0, n = 21 and 20 plants, respectively) to the expected phenotype class. Groups of mutant plants transformed with 3xHA-AGO7 constructs were predominantly assigned to one or the other class: primary transformants for 422 bp to 1934 bp promoter constructs were mostly scored as complemented, whereas most transformants for 298 bp and 0 bp promoter construct displayed the downward-curled-leaf mutant defect (Figure 6).

**Figure 6.**
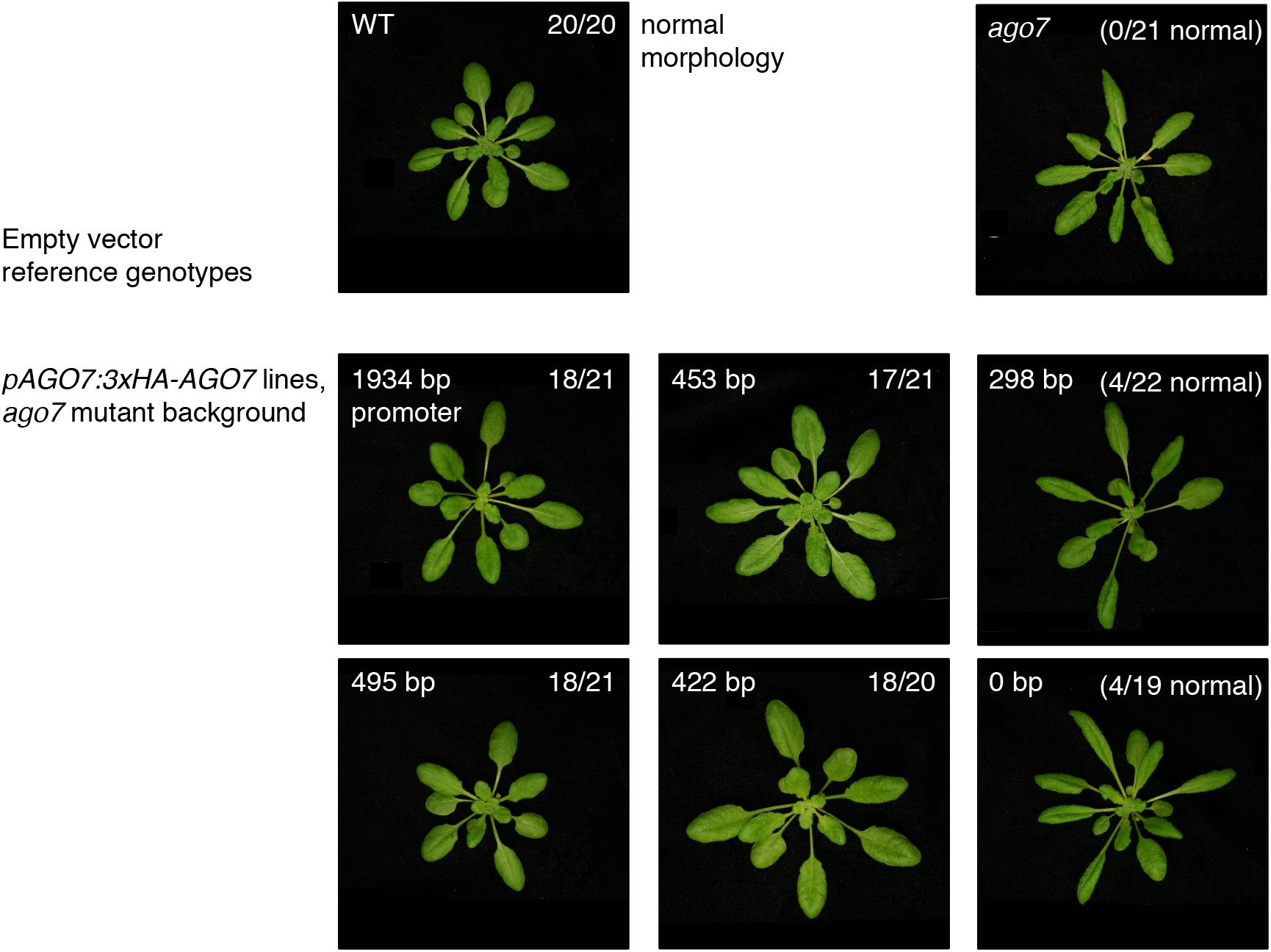
Complementation of *ago7* mutant leaf shape phenotype (top right) with 3xHA-AGO7 transgenes driven by truncated versions of the *AGO7* promoter. One representative (major class) primary transformant is shown for each genotype. Upper-left corner labels for middle and bottom rows indicate the length of upstream *AGO7* regulatory sequence used to drive the 3xHA-AGO7 coding sequence in each construct. Upper-right corner numbers indicate the fraction of plants blindly assigned to the normal morphology category.

**Figure 7.**
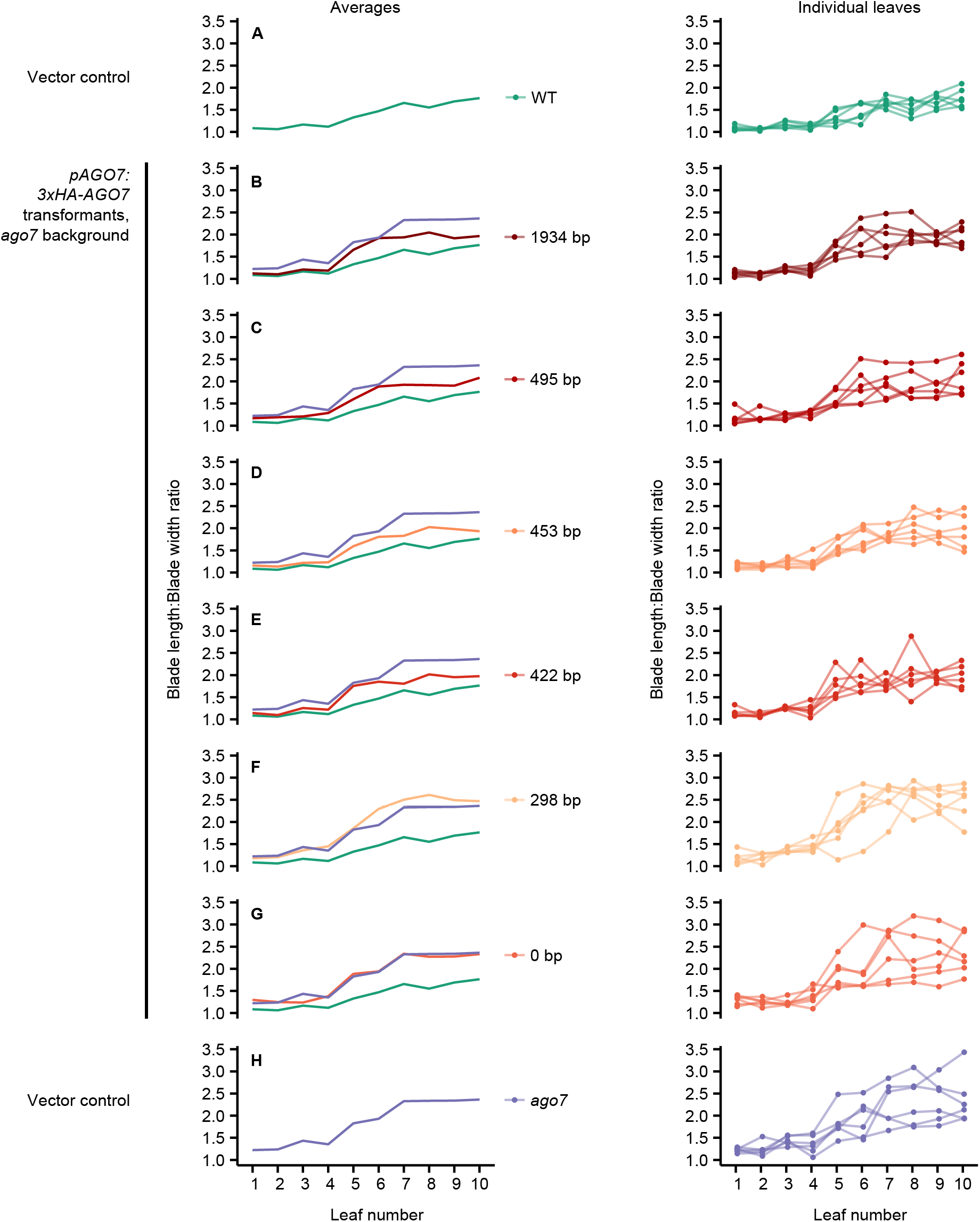
Transgenic complementation of *ago7* leaf shape defects, quantified based on leaf blade length to width ratio for true leaves 1 to 10. Values for each individual plant are connected with lines on the right-hand graphs, and the average of these values is plotted on the left. Averages for empty vector control genotypes (panels A and H) are repeated in each left-hand panel to facilitate comparison.

We extended this result by quantifying leaf shape for a smaller number of transformants, by dissecting, scanning, and measuring leaves in order [50]. For the reference genotypes, leaf blade length-to-width ratios were higher for wild-type relative to mutant plants, due to increased curling and/or elongation (Figure 7, panels A and H). Promoterless and 298 bp promoter construct transformants were not distinguishable from empty vector mutant controls (Figure 7, panels F and G). Longer promoter constructs shifted blade length-to-width ratios down towards wild-type levels (Figure 7, panels B to E), which we interpret as partial complementation, consistent with the rosette-level results in Figure 6. Independently measuring these leaf dimensions at one position (true leaf 6) with calipers yielded similar results (Figure S5).

Results were similar for a related metric that quantifies leaf elongation, the ratio of leaf blade length to petiole length (Figures S5 and S6). The difference between wild-type and mutant background control plants was smaller for this metric (Figure S6, panels A and H), as was the difference, if any, between means for the 1934 bp promoter construct lines and wild-type empty vector control lines (Figure S6B). Means were longer at most leaf positions (i.e. closer to wild-type) for intermediate-length promoter constructs (Figure S6, panels C, D, and E) than for short promoter constructs (Figure S6, panels F and G). Exceptions at one position (true leaf 10) were caused by recently emerged leaf “outliers”, the petioles of which were very short and thus disproportionately affected by technical variation (Figure S6C).

The promoter lengths tested end immediately adjacent to core SPL and TCP binding sites (two ‘TGGTCC’ sites and one of two ‘GTAC’ motifs discussed above; see Figure S7). The 422 bp promoter transgene lacks all of these sites, but is sufficient for partial complementation (Figure 6, Figure 7E, Figure S6E). We therefore tentatively conclude that SPL and TCP binding is not required for *AGO7* transcription at levels that are sufficient for normal leaf morphology. The morphological data described allow us to estimate possible small differences between leaf shape in the complemented lines, but further experimentation would be necessary to relate such differences to cellular parameters or promoter structure.

Finally, we scored appearance on trichomes on abaxial leaf surfaces to assess complementation of the forward shift in *ago7* mutants [3]. Consistent with results from previous transgenic experiments [13, 51], abaxial trichomes were visible on an earlier leaf for empty-vector-transformed mutant plants relative to corresponding wild-type plants (Figure S7); abaxial trichomes appeared 1.7 leaf positions earlier on average (95% confidence interval 0.5 to 2.9, *p* = 4 × 10^−4^, Tukey’s honest significant difference method). However, there was considerable variability, possibly due to effects from hygromycin selection. No 3xHA-AGO7 transgenic line showed a detectable increase in earliest abaxial trichome position (relative to empty-vector-transformed mutant plants; p > 0.3), indicating that none of the promoter lengths tested were able to drive full complementation of this defect. Alternative strategies may be required to assess ARF-mediated effects of AGO7 levels on trichome production.

## 3 Discussion

We characterized the structure of the *AGO7* promoter with transgenic analyses and a large-scale screen for upstream regulators. Figure 8 provides a possible interpretation these results in terms of TF binding events. The most notable result from our Y1H analysis was a direct connection to multiple miR156-targeted SPL and miR319-targeted TCP factors. This result appears to reinforce the idea that gradual repression of *MIR156* transcription is the key regulatory step controlling heteroblasty in plants [40]. This connection, if verified in future studies, provides an additional example of functional linkage between SPL and TCP TFs [52–54], and a link to ARF repressors, the other main regulators of dynamic changes in leaf shape (Figure 9). However, we were not able to assign a clear function to the candidate SPL and TCP binding sites in the *AGO7* promoter, particularly because a 422 bp proximal promoter region lacking all these sites is sufficient for substantial transgenic complementation of leaf morphology defects in *ago7* mutants (Figures 6, 7, S5, and S6).

**Figure 8.**
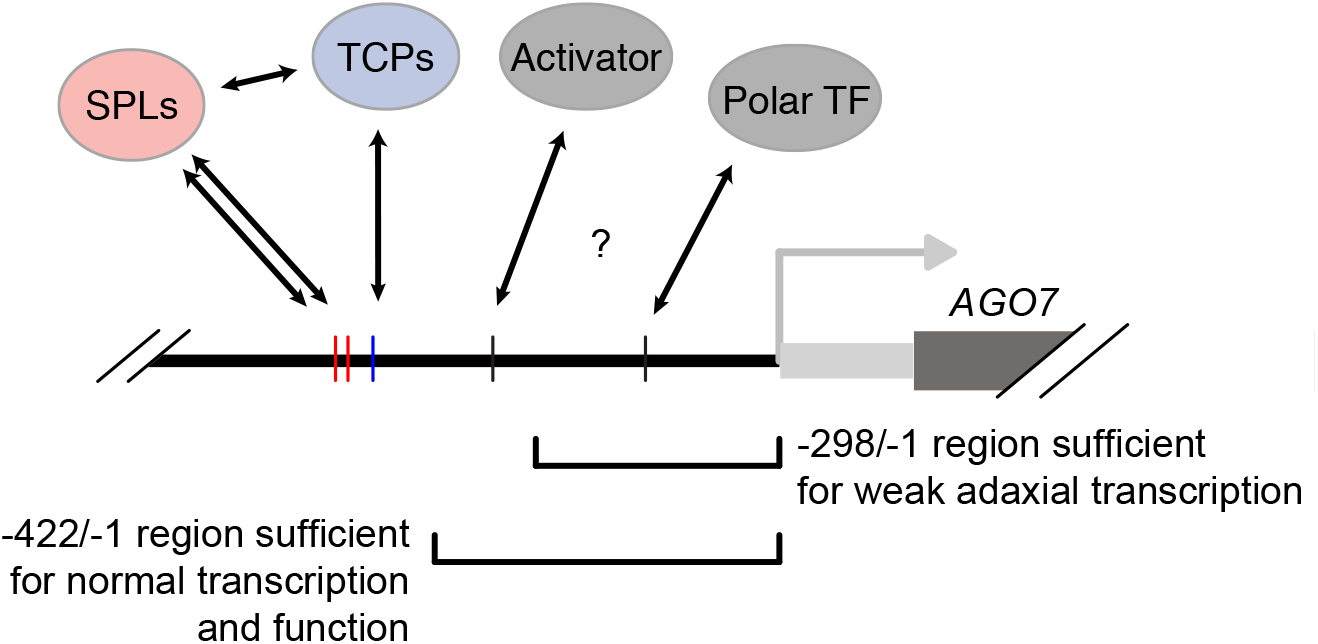
Schematic of the *AGO7* proximal promoter region with hypothesized TF binding sites and summary results from transgenic analyses indicated. One or more activator and polarity determinant TFs are proposed to bind at undetermined sites in the regions indicated, as discussed in the text.

**Figure 9.**
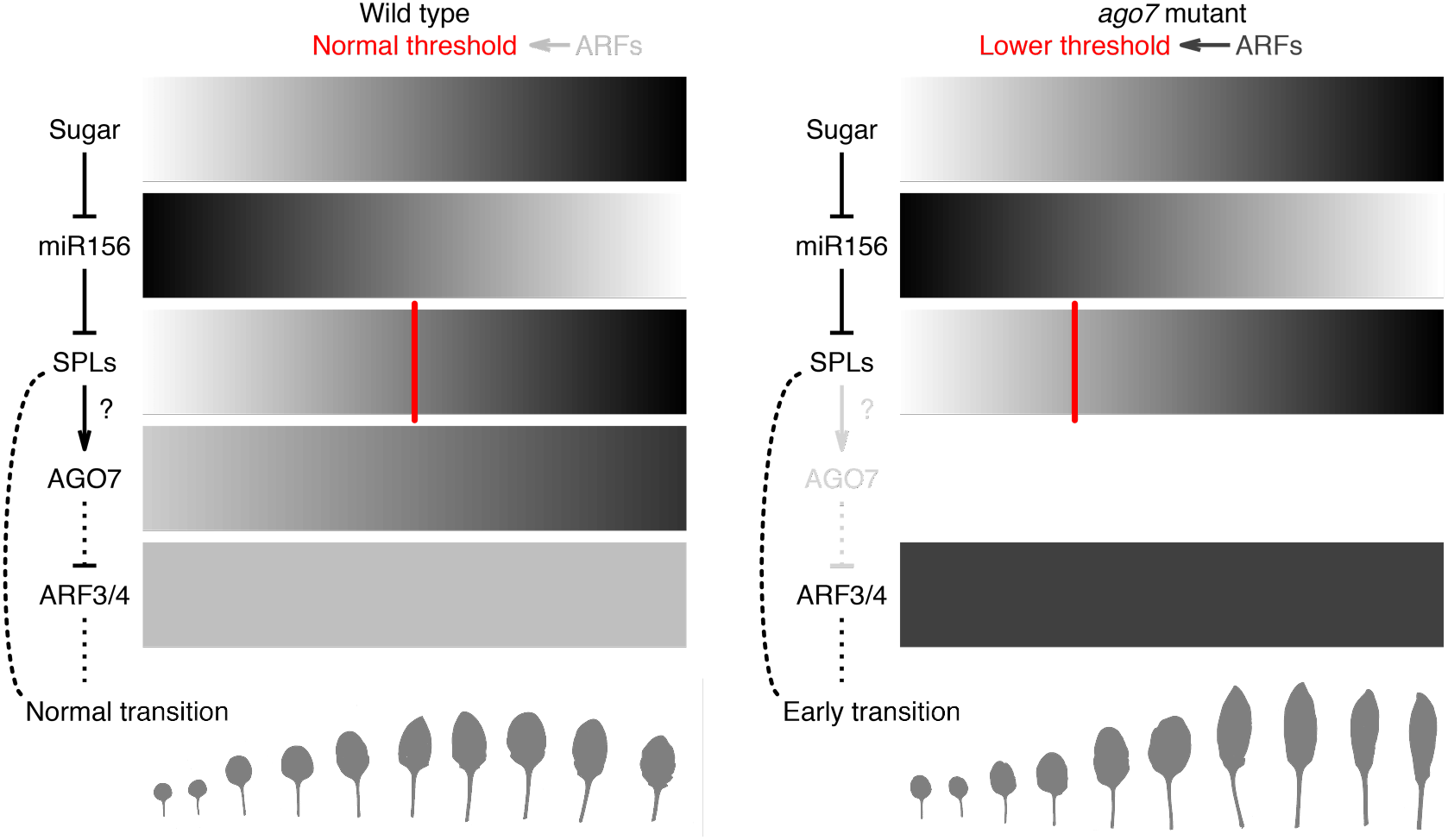
Revised model for control of heteroblasty in *A. thaliana*, incorporating the likely activation of *AGO7* by SPLs. SPL levels gradually increase until they reach a hypothetical activity threshold, controlled in part by ARFs, and trigger changes in leaf characteristics [20, 55]. When the AGO7 pathway is disrupted, ARF levels go up, lowering the threshold for transition such that it is reached earlier. This perturbation results in leaves that are thinner, longer, and more curled in *ago7* mutants. Indirect repression of ARFs via AGO7 and *TAS3* tasiRNAs may have significance for feedback control of SPL activity.

Our truncation analysis provided preliminary evidence for two other functional regions of the *AGO7* promoter (Figure 8). We obtained different outcomes for mutant plants tested with 422 bp promoter constructs (largely complemented) versus 298 bp promoter constructs (not complemented). This difference suggests that one or more functionally important binding sites is present in the −422/−299 region. In general agreement with this idea, signal was qualitatively weaker for a 298 bp promoter:GUS reporter than for the next-longest promoter fragment tested (Figure 5). Multiple experiments suggest that the minimal core promoter and possibly one or more polarizing *cis* elements are intact in the 298 bp proximal region, but dissecting this further has been technically challenging because of the faintness of the signal.

Despite progress, we did not succeed in identifying TF binding events necessary and/or sufficient for polar expression of *AGO7* and *AGO10*. The *YABBY1/FILAMENTOUS FLOWER* gene is one promising candidate because of its well-known role in polarity [56, 57]: as documented in the supplemental materials [35], YAB1 is predicted to have high affinity for a site in the *AGO7* proximal promoter but did not emerge as a hit from the Y1H screens. Surprisingly, we also did not recover the polarity factor REVOLUTA for the *AGO10* promoter [58]; this likely represents a biological false negative. *AGO1* is ubiquitously expressed [11], and therefore expected to be under very robust transcriptional control which may be difficult to dissect. The Y1H results presented here should be a useful resource as further genome-wide chromatin immunoprecipitation data for *A. thaliana* become available.

The truncation strategy used for our transgenic assays preserves the distance between *cis* elements, but also has inherent limitations. We did not test the possibility that SPL and TCP core binding sites are *sufficient* for specific genetic functions. The apparent enhancer(s) in the −422/−299 region may be functionally redundant with these binding sites, and may therefore have masked any contributions to morphology through *AGO7*. Redundant clusters of activator binding sites are thought to be common, and may contribute to robustness [59–62]. Effects may be larger in other tissues, given the important functions of ARF repressors in fruits and roots [63–67]. Alternatively, the sites may simply be nonfunctional, at least in *A. thaliana*. Testing SPL and TCP binding in multiple tissues would help in assessing these possibilities; such data would aid evaluation of the possibility that the *AGO7* promoter integrates both temporal and spatial signals. More broadly, linking such TF binding events to changes at the cellular level in diverse plants should remain a challenging but productive approach [68, 69].

## 4 Methods

### 4.1 Plasmid construction

Promoter fragments were PCR-amplified from previously described plasmids [13], with the primers listed in Table 1. Truncated and modified forms of the *AGO7* promoter were made with the oligonucleotides listed in Table 2 and Table 3. Gel-purified PCR products were cloned with the pENTR D-TOPO kit (Invitrogen) and LR-recombined into several destination vectors: pGLacZi for Y1H screens [70], pMDC162 for GUS transcriptional reporters, and pMDC99 for transgenic complementation assays [71]. The destination vector pY1-gLUC59(GW) used for the secreted *Gaussia* luciferase Y1H reporter system has been described [41].

**Table 1.**
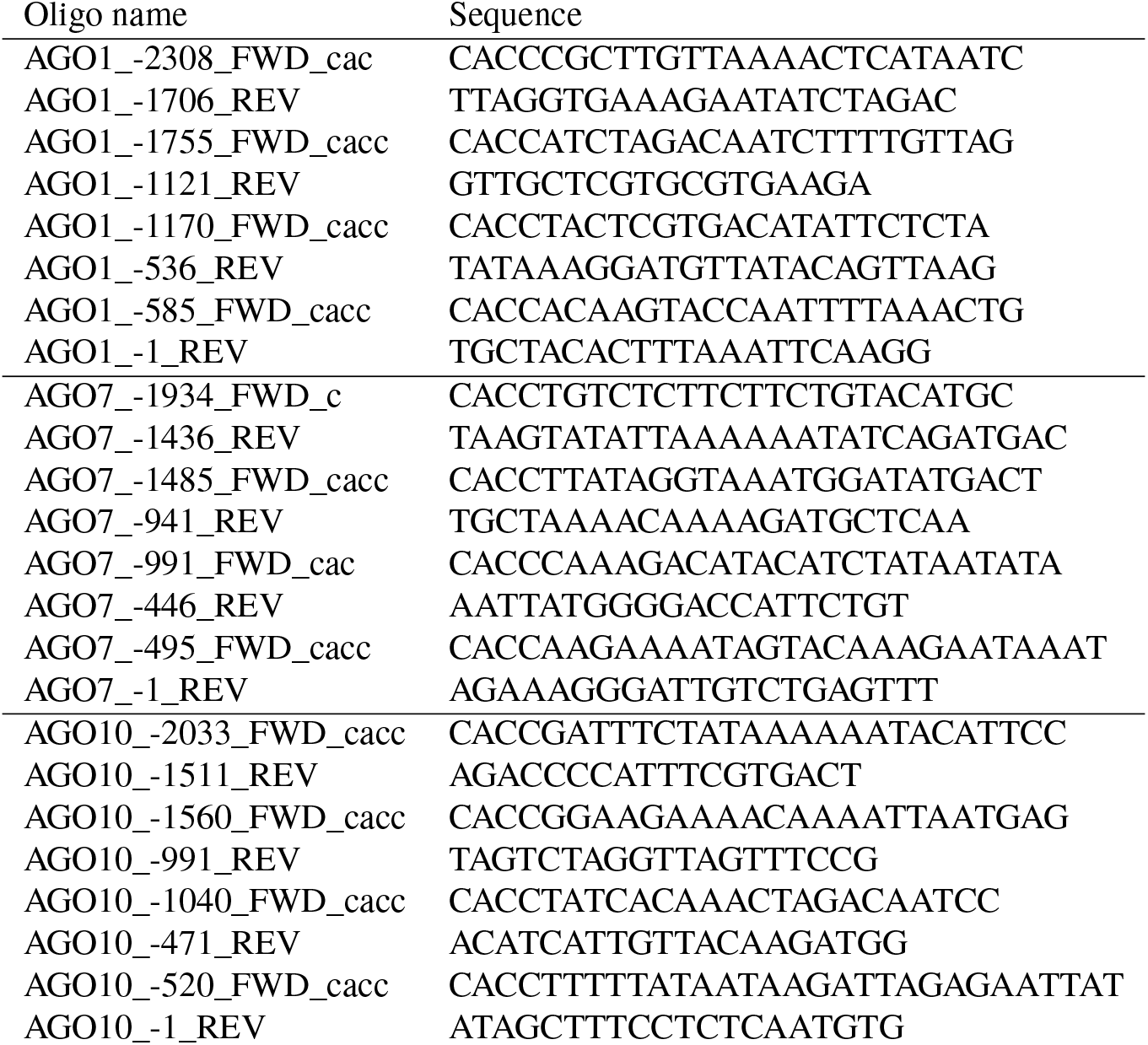
Oligonucleotide sequences used for *AGO* promoter TOPO cloning. Primer names indicate position of 5’-most genomic base relative to the annotated transcription start site. Names also list the nucleotides added to create ‘CACC’ sequences for directional TOPO cloning.

**Table 2.**
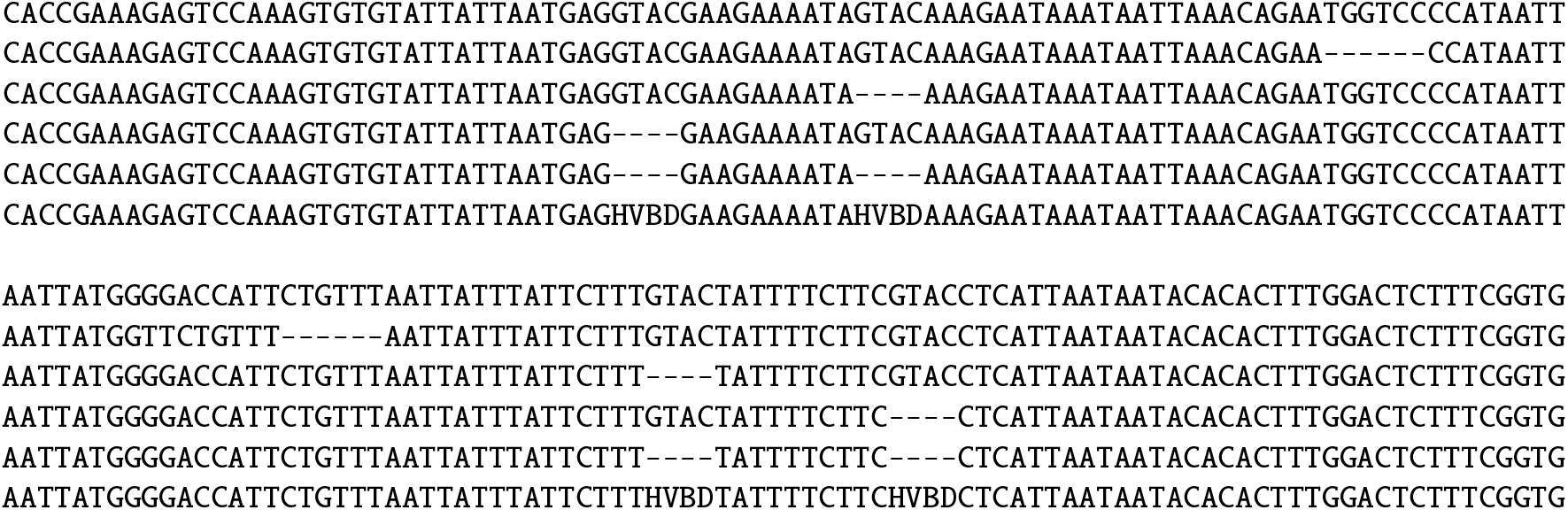
Oligonucleotide sequences directly cloned for *AGO7* promoter mutation analysis in yeast. Forward sequences (5’ to 3’ in the direction of *AGO7* transcription) are followed by corresponding reverse sequences. Dashes indicate bases “deleted” relative to the genomic reference sequence. The last sequence for each set is a degenerate oligo; a single clone resulting from these oligos was used, as indicated in the caption for Figure 2.

**Table 3.**
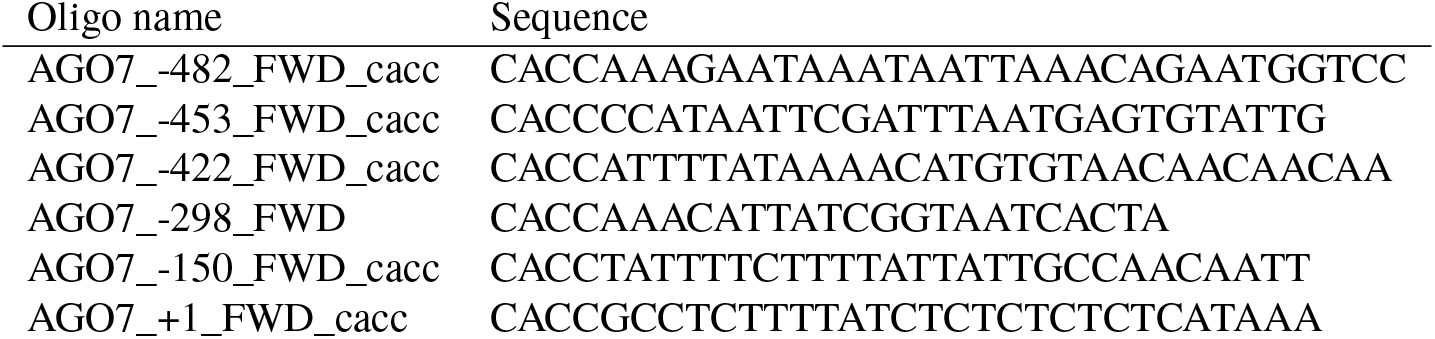
Forward primer sequences used for TOPO cloning of truncated versions of the *AGO7* promoter. Names follow Table 1. Bases −298/−295 are a natural ‘CACC’ sequence suitable for directional TOPO cloning.

### 4.2 Y1H screens

Automated *lacZ* screens were done as previously described [31, 32] using a collection of 1497 TFs and an Agilent BioCel 1200 robotic platform. The TF-activation domain fusion yeast strain collection (arrayed in 384-well plates) was mated to bait strains. Diploid cells were selected in media lacking uracil and tryptophan, lysed by freeze-thaw, and assayed for *β*-galactosidase activity. Targeted Y1H assays were done similarly, with the lysis and assay steps replaced, essentially as described [41]. Briefly, diploid cells were resuspended in phosphate-buffer saline, 50 μL of cells were transferred to a clear-bottom plate, and a Synergy H1 plate reader (Biotek) was used to inject 10 μL of 20 μM coelenterazine substrate solution into each well and read luminescence immediately afterward (0.1 s integration time).

### 4.3 Plant materials and growth conditions

All *A. thaliana* plants descended from the reference Col-0 accession. The *zippy-1* mutant allele was isolated by Hunter et al. [3], and is referred to throughout as “*ago7*”. Plants were transformed by floral dip using *Agrobacterium* strain GV3101 [72, 73].

Plants were grown under short day conditions (8 hours light, 16 hours dark) in a Conviron MTR25 reach-in chamber with PolyLux fluorescent bulbs (200 μmol photons per second per square meter) at 22 °C with 50% humidity.

### 4.4 *ago7* mutant complementation tests

Measurement of leaf phenotypes followed previous work [18]: we scored the index of the earliest leaf with at least one abaxial trichome using a stereomicroscope at 28 to 30 days post-stratification, and concurrently measured the blade length, blade length, and petiole length for the sixth true leaf with digital calipers (Mitutoyo, Japan). At a later timepoint (33 and 35 days post-stratification), we dissected and scanned the first ten true leaves from each plant with a Canon Pixma MP190 flatbed scanner. Leaf shape parameters were measured with the LeafJ plug-in for ImageJ [50]. Plants were also photographed from above from 11 days post-stratification onward and the time-lapse image data [74] documented with the rest of the experiment [75–78].

### 4.5 GUS assays

Histological GUS assays were essentially as described [12, 79, 80]. Seedlings were collected into ice-cold 90% acetone, incubated at −20 °C for 20 minutes and then room temperature for another 20 minutes. Seedlings were washed twice (5 minutes each) with staining buffer (100 mM sodium phosphate [pH 7], 20% methanol, 0.1% Triton X-100, 1.5 mM ferri- and ferrocyanide).

Staining buffer with 0.5 mg/mL 5-bromo-4-chloro-3-indolyl-*β*-d-glucuronic acid (X-Gluc) was vacuum-infiltrated into seedlings on ice for two rounds of 15 minutes each. Samples were then incubated at 37 °C for 20 hours, taken through an ethanol/histoclear series, and infiltrated with Paraplast Plus at 60 °C, before embedding [79]. Tissue sections (10 μm thickness) were mounted on Probe-On Plus slides (Thermo Fisher), deparaffinized with histoclear, and coverslipped. Sections were viewed and photographed with a Leica DM750 microscope and ICC50 HD camera.

### 4.6 Data and code availability

As noted above, data and software code supporting this manuscript have been deposited as Zenodo records 1258642, 1322798, 1340636, 1345229, and 1472235 [35, 36, 49, 75, 76]. Data processing was done with the R Statistical Computing Environment [81] and the Bioconductor BioStrings package was used for PWM scans [82, 83]. The relevant data supplement [35] also includes results from ‘Find Individual Occurences of Motifs’ tool (FIMO) scans [45] done via the online MEME Suite [84] version 4.12.0, with default settings (*p* < 10^−4^ cutoff) and three collections of DNA-binding specificity models [43, 46, 47].

## Acknowledgments

We thank Danforth Center Plant Growth Facility staff for excellent plant care, G. Nguyen and R. Allscheid for logistical support, and members of the Carrington lab for helpful discussions. D.H. Chitwood and J.G. Hodge provided essential advice on histological analysis. We thank T.C. Mockler and members of his lab (J. Gierer, D. O’Brien, M. Wiechert) for assistance with their plate readers and liquid-handling equipment. This work was supported by US National Science Foundation award 1330562 (to JCC) and US National Institute of Health grants AI043288 (to JCC), GM056006 (to SAK and JLP-P), and GM067837 (to SAK). JSH was supported by an NSF graduate research fellowship (award 1143954).

## Author contributions

JSH, JLP-P, GB, SAK, and JCC designed the research. JSH, JLP-P, GB, MAH, EEH, HF, KMB and JM performed research. JSH, JLP-P, MAH, EEH, KMB, and JCC analyzed data. JSH and JCC drafted the paper; all authors commented on and approved the paper.

---

**Figure S1.**
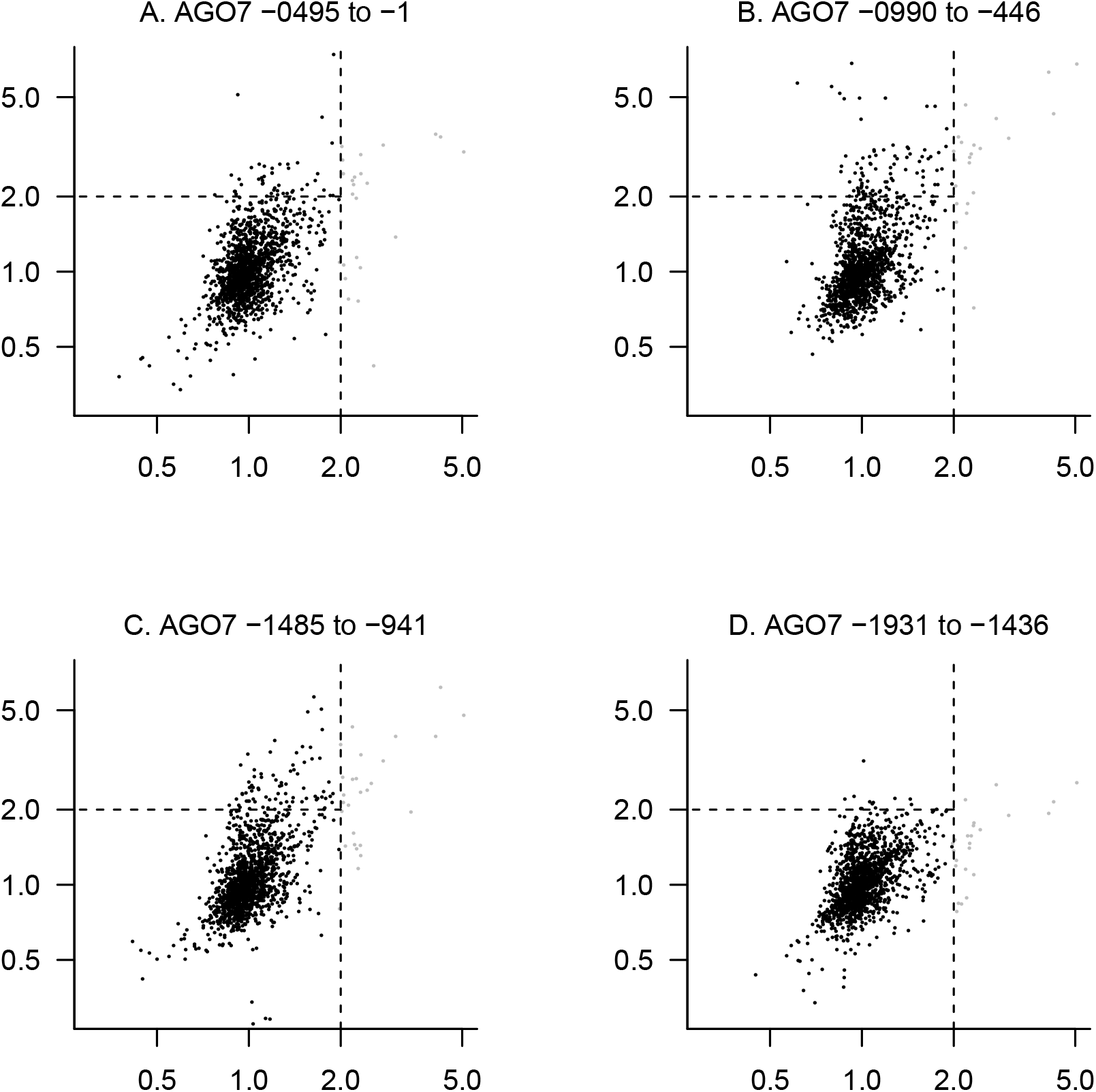
Scatterplots of *β*-gal activities with likely nonspecific activatiors indicated for *AGO7* promoter fragment screens. Panel B is equivalent to Figure 1C.

**Figure S2.**
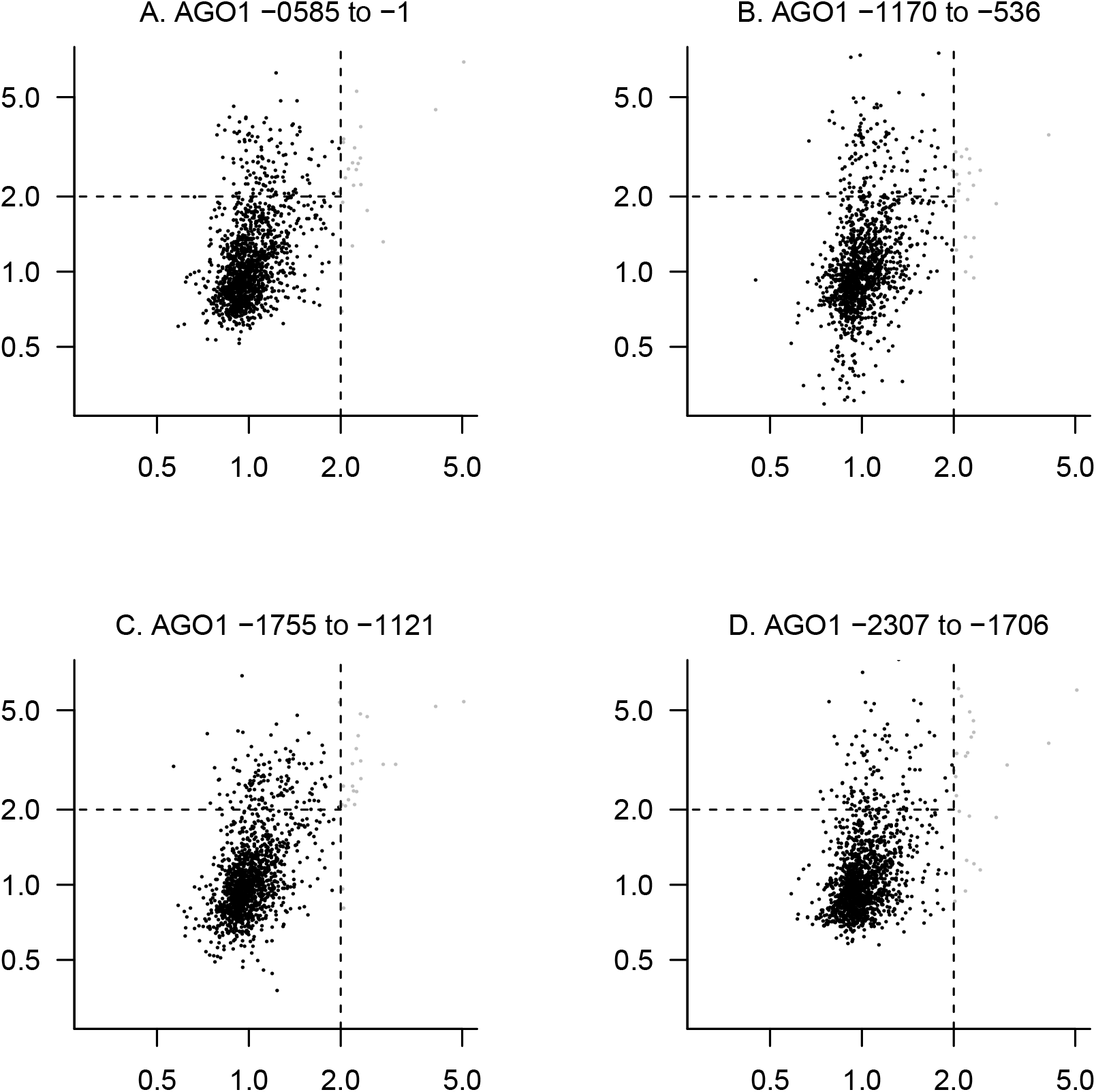
Scatterplots of *β*-gal activities with likely nonspecific activatiors indicated as in Figure 1C for *AGO1* promoter fragment screens.

**Figure S3.**
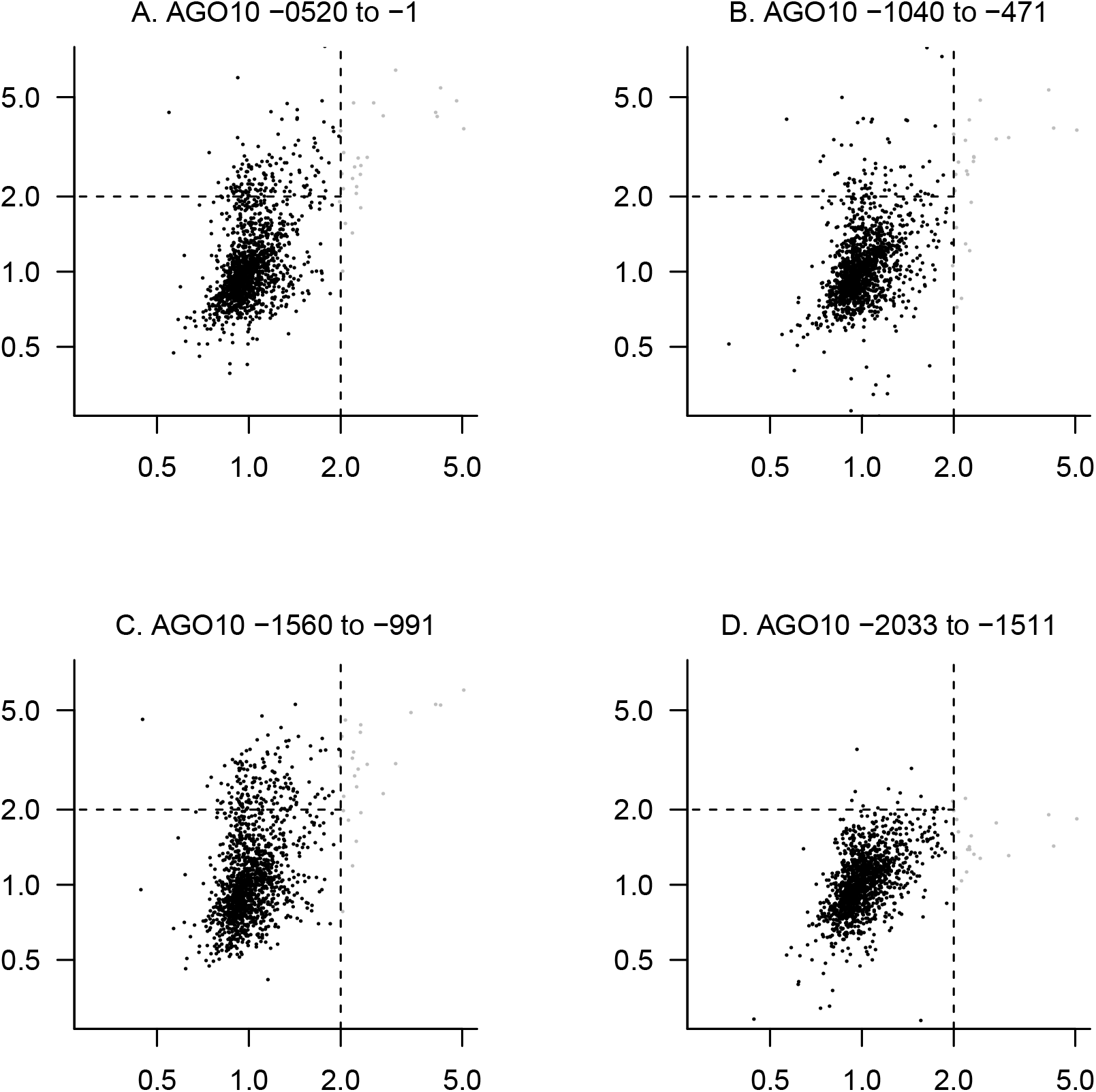
Scatterplots of *β*-gal activities with likely nonspecific activatiors indicated as in Figure 1C for *AGO10* promoter fragment screens.

**Figure S4.**
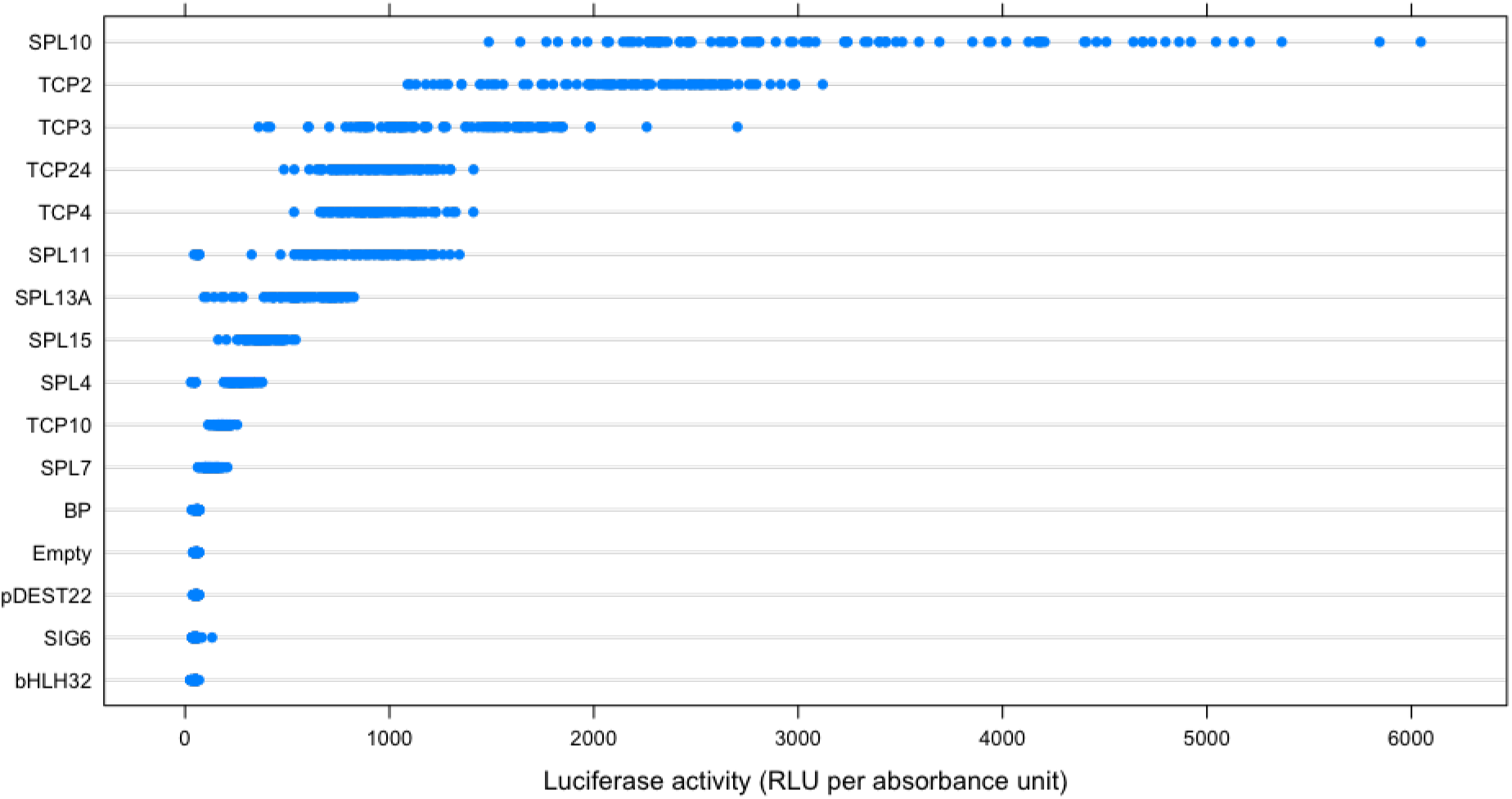
Targeted Y1H assay using *Gaussia* luciferase reporter, quantified in terms of relative luminescence units per absorbance unit at 600 nm. TFs were tested against the *AGO7* −990/−446 region and are displayed in order by mean reporter activity.

**Figure S5.**
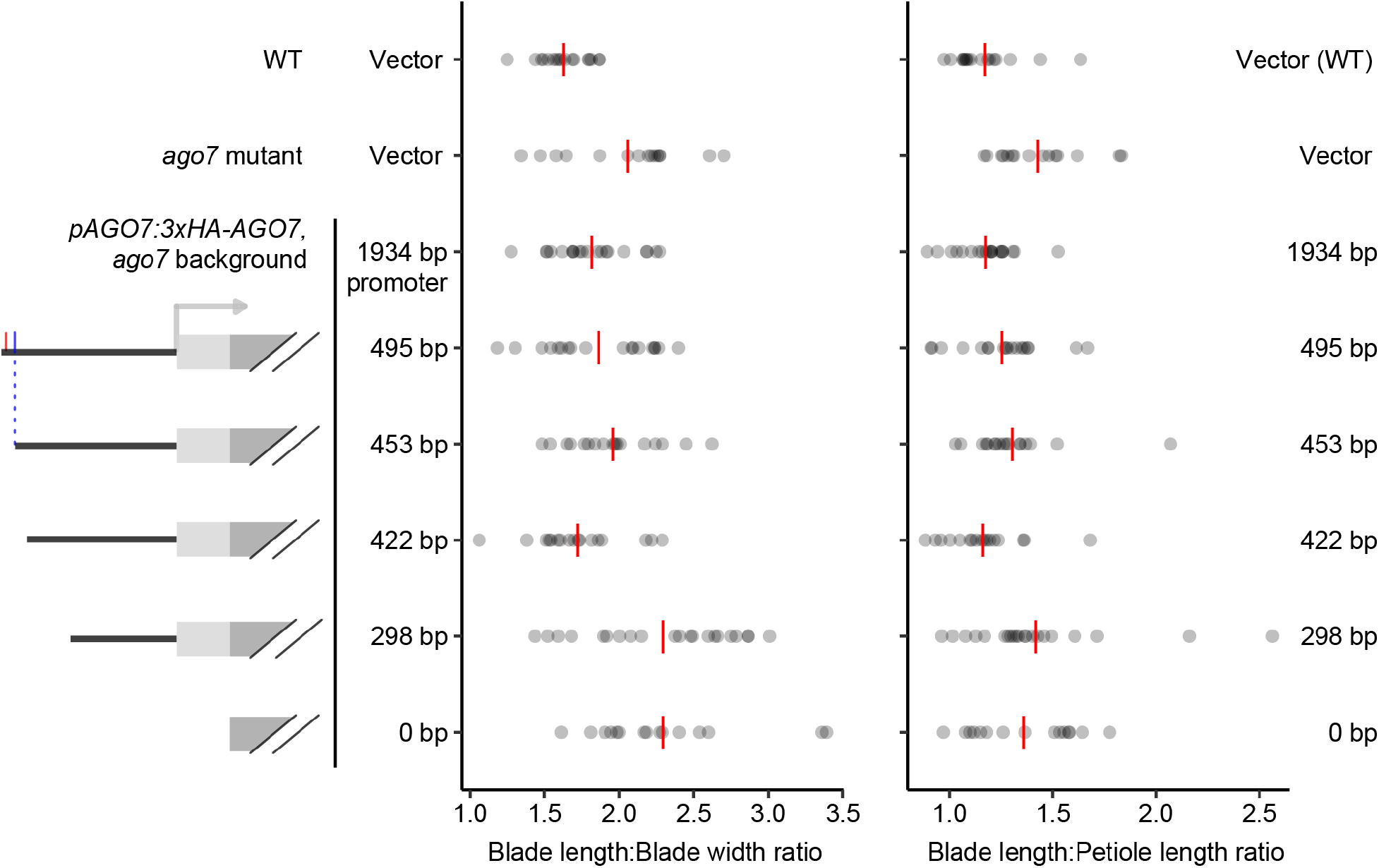
Complementation of *ago7* leaf shape defects, quantified based on leaf blade length-to-width ratio (left) and leaf-blade-length to petiole-length ratio (right) measured for true leaf 6 with calipers on days 28 to 30 days post-stratification. Each datapoint shows the ratio for a distinct primary transformant. Red lines indicate the mean for each genotype.

**Figure S6.**
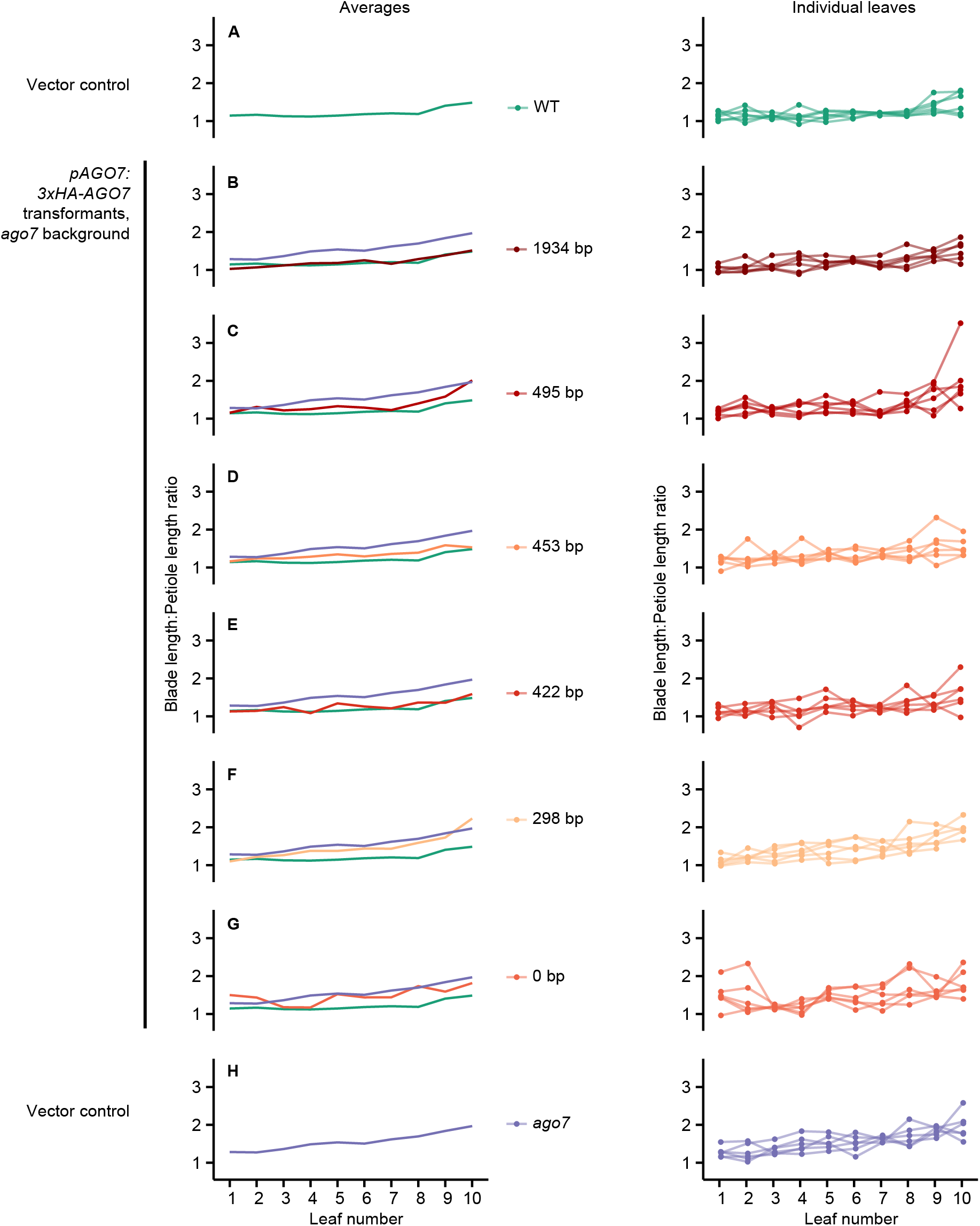
Complementation of *ago7* leaf shape defects, quantified based on leaf blade length to petiole length ratio. Panel layout is as in Figure 7.

**Figure S7.**
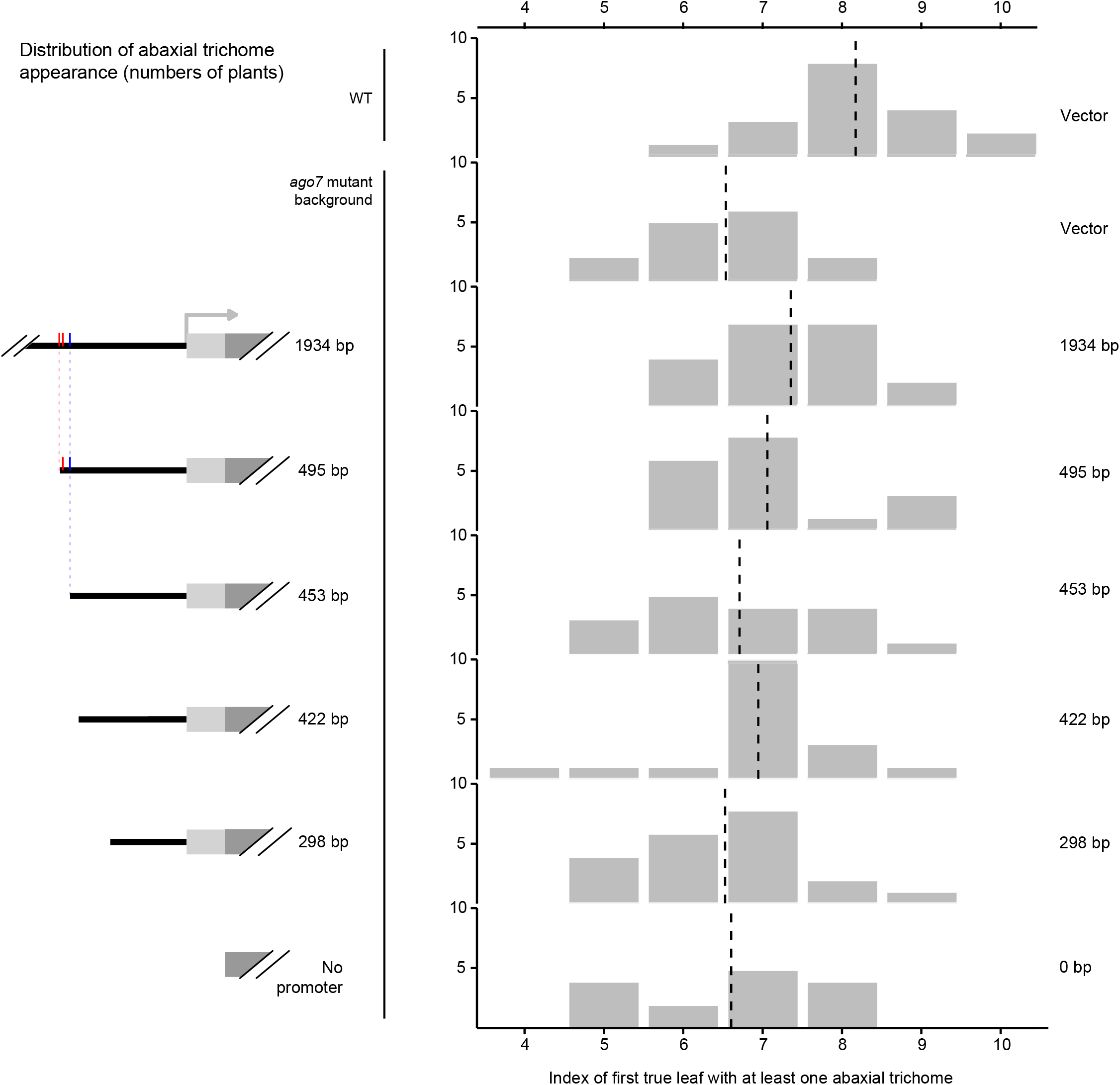
Assay for complementation of *ago7* early abaxial trichome appearance phenotype with 3xHA-AGO7 transgenes driven by truncated versions of the *AGO7* promoter. Dashed lines indicate the mean for each genotype. Core SPL (red) and TCP (blue) binding sites are indicated with tick marks.

